# Development of male-sterile lines of *Setaria viridis* to accelerate C_4_ model plant genetics

**DOI:** 10.1101/2025.07.14.664229

**Authors:** Hui Jiang, Erik A Myers, Britney Millman, Dong-Yeon Lee, Dmitri A. Nusinow, Daniel F Voytas, Ivan Baxter

## Abstract

*Setaria viridis* is a diploid C_4_ grass in the Poaceae family, notable for its rapid life cycle of 6–8 weeks from sowing to seed—much shorter than the 4–5 months required by crops such as *Zea mays* and *Sorghum bicolor*. This fast growth makes *S. viridis* a valuable model for C_4_ crop research. Genetic crosses are essential for studying gene function, but manual crossing is labor-intensive and time-consuming. To address this, we developed a male-sterile line by targeting the *S. viridis* ortholog of *Setaria italica* NO POLLEN 1 (*SiNP1*), which encodes a glucose–methanol–choline oxidoreductase required for pollen exine formation. Using Cas9 and TREX2-mediated genome editing, we generated *SiNP1* knockouts in both the *S. viridis* ME034V and A10.1 backgrounds that were fully male-sterile. Backcrossing T_0_ male-sterile plants to ME034V yielded a stable line homozygous for a 59 bp deletion, easily genotyped by PCR. Using this line, we developed a simple and efficient crossing protocol that eliminates the need for emasculation. This method enables a single person to perform up to 100 crosses per day—compared to 15 using traditional methods—and yields 20–32 F_1_ hybrid seeds per panicle with 100% genetic purity. We also quantified pollen flow and outcrossing frequencies under greenhouse conditions to develop optimal bagging strategies and prevent unintended pollination.This resource accelerates genetic research in *S. viridis*, enhancing its utility as a premier C_4_ model for mapping and functional genomics.

## Introduction

*Setaria viridis* (*S. viridis*) is a C_4_ grass in the Poaceae family, which includes agronomically important crops, such as maize, sorghum, wheat, barley, and rice. Compared to other Poaceae, *S. viridis* has a rapid life cycle, progressing from seed to seed in only 6 to 8 weeks (Sebastian et al. 2014). In addition, *S. viridis* has a well-annotated publicly available genome, and accessions A10.1 and ME034V are readily transformable via Agrobacterium (Mamidi et al. 2020; Brutnell et al. 2010; Nguyen et al. 2020). Recent advancements have established key genetic resources and tools for both forward and reverse genetics. Two NMU mutagenized populations (Huang et al. 2017) (Chatterjee et al. 2021; Coe et al. 2018) have enabled mapping of genes associated with agronomically important traits such as inflorescence architecture (Huang et al. 2017); (Yang et al. 2018; Yang et al. 2021), crown root development (Sebastian et al. 2016), and C_4_ photosynthesis (Chatterjee et al. 2021). Resequencing of 598 *S. viridis* accessions has further provided a foundation for genome-wide association studies (GWAS) across a broad range of traits (Mamidi et al. 2020). As such, *S. viridis* has emerged as a powerful model species, enabling discoveries that can be translated to other members of the Poaceae.

Despite these advances, the self-pollinating nature of *S. viridis* presents challenges for generating genetic crosses. The flowers are small, densely packed, and exhibit a narrow (∼40-minute) anthesis window (Supplementary Figure 1). Current crossing protocols (Jiang et al. 2013) rely on heat emasculation, This labor-intensive method requires considerable expertise and often yields variable success across different accessions, necessitating genotype-specific optimization (Jiang et al. 2013).

Male sterility is a key breeding tool that facilitates genetic crosses in many plant species. It is broadly categorized into Cytoplasmic Male Sterility (CMS) or Genic Male Sterility (GMS). CMS arises from interspecific crosses and results in progeny with a nuclear genome from one parent and a mitochondrial genome from the other (Toriyama 2021). Male sterility in CMS lines is caused by a defective mitochondrial gene and a nuclear genome that lacks the corresponding restorer of fertility (*Rf*) gene (Schnable 1998). A CMS system typically requires three-lines: a male-sterile line (with defective cytoplasm and no *Rf* gene), a maintainer line (normal cytoplasm but lacking the *Rf* gene) and a restorer line (carrying the *Rf* gene and desired agronomic traits).

In contrast, GMS results from a mutation in a single nuclear gene. GMS systems require only two lines: a male-sterile line and a maintainer line, both derived from the same genetic background. Maintenance of the male-sterile line is achieved by self-fertilization of heterozygous plants. Mendelian segregation produces progeny in which approximately 25% of seeds are male-sterile. To date, no male-sterile lines have been reported for *S. viridis*.

In *S. italica*, an ethyl methanesulfonate (EMS) mutant screen identified *NO POLLEN 1* (*SiNP1, Seita*.*9G347800*) as essential for male fertility (Zhang et al. 2021). *SiNP1* is orthologous to male sterility genes in maize, rice, and Arabidopsis (Liu et al. 2017; Chen et al. 2017; Krolikowski et al. 2003). In both *S. italica* and maize, *NP1* encodes a glucose-methanol-choline oxidoreductase (GMCO) that is predominantly expressed in the tapetal cells of the anther (Chen et al. 2017; Zhang et al. 2021). The tapetum contains two key organelles, elaioplasts and tapetosomes. Tapetosomes, derived from the endoplasmic reticulum (ER), encase lipid droplets that form by budding from the ER. During pollen development, tapetal cells undergo programmed cell death, releasing the contents of the tapetosomes onto the pollen surface (Hsieh & Huang 2007). The tapetum also synthesizes sporopollenin precursors that contribute to formation of the pollen exine, a protective outer layer (Lallemand et al. 2013). Disruption of *SiNP1* causes abnormal exine formation and results in non-viable pollen (Zhang et al. 2021).

Here, we employed a robust gene-editing approach to target the *SiNP1* ortholog in *S. viridis*. Using Cas9 and TREX2 (a 3’ exonuclease), we generated homozygous deletions in *SvNP1* in the T_0_ generation, yielding fully male-sterile mutants. Leveraging this line, we developed a simple and highly efficient crossing protocol for *S. viridis* that eliminates the need for emasculation. These mutants enabled us to assess pollen flow in greenhouse-grown *S. viridis* and offer a valuable tool to streamline genetic crosses in this model species.

## Results

### Robust mutagenesis of *SvNP1*

The *S. viridis* homolog of *SiNP1* comprises four exons. To generate deletions in the coding sequence via CRISPR/Cas9-mediated mutagenesis, we designed pairs of single guide RNAs (sgRNAs) targeting exon 2 or 3 of *SvNP1* (Figure 1A). The distance between sgRNA target sites ranged from ∼100 to 400 bp. Each sgRNA was expressed from the *OsU6* promoter to avoid the RNA silencing issues associated with the pol II *CmYLCV* promoter (Weiss et al. 2020). Further, *OsU6* has shown high efficiency for heritable, multiplexed gene editing in rice (Wu et al. 2024).

**Figure 1.**
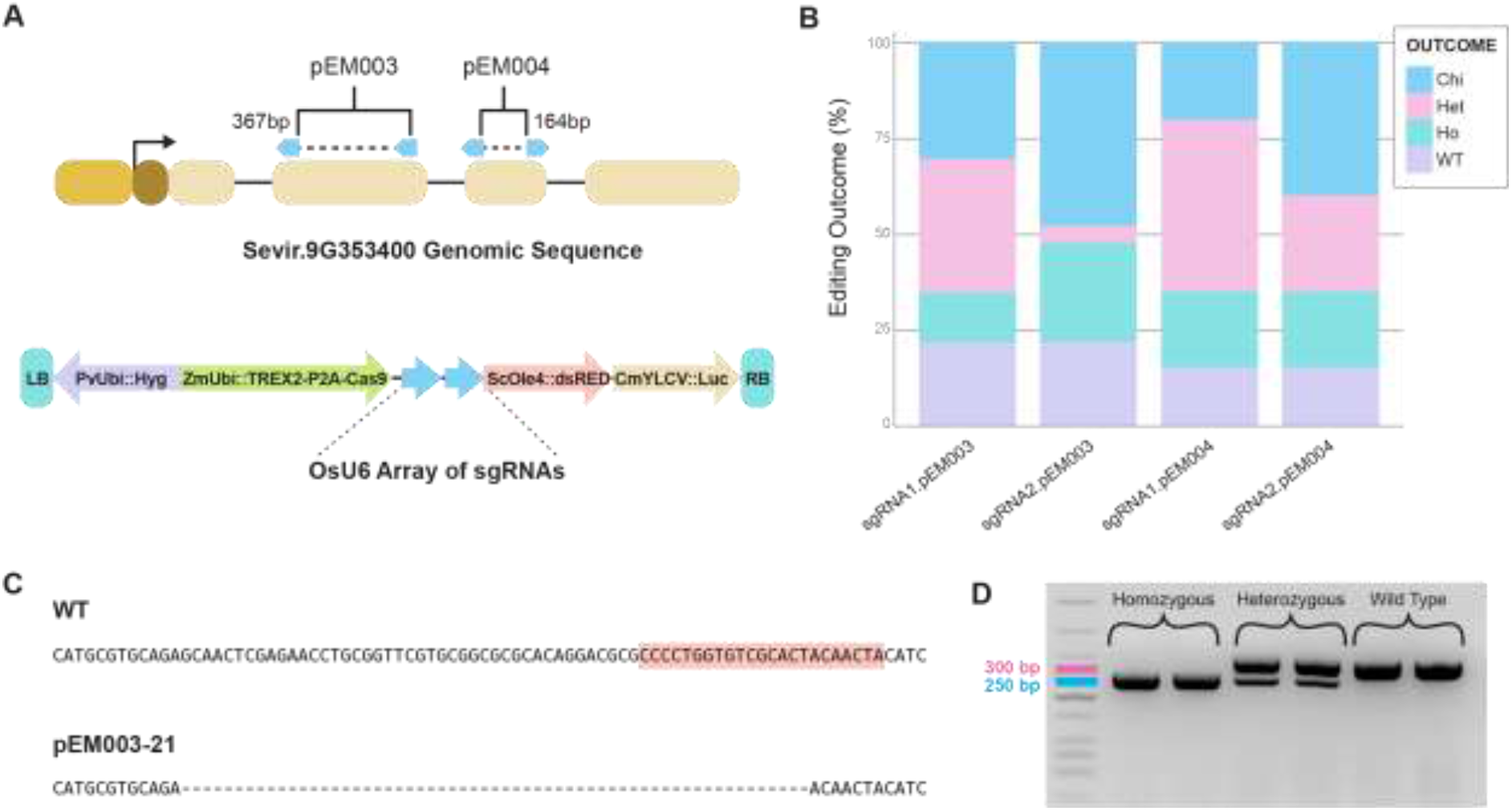
CRISPR/Cas9 gene-editing of *SvNP1*. **A)** The top schematic represents the *SvNP1* genomic locus, and blue arrows denote sgRNA target sites. pEM003 and pEM004 target exons 2 and 3, respectively. The bottom schematic represents the T-DNA used to deliver the gene-editing reagents through Agrobacterium-mediated transformation of *S. viridis*. Details of the gene constructs are described in the text. **B)** The relative proportion of different editing outcomes at each sgRNA target site for both pEM003 and pEM004 in T_0_ plants. A heterozygous (Het) editing outcome means that the plant has different alleles on each chromosome; homozygous (Ho) refers to identical, non-wild-type alleles on both chromosomes. If editing occurred, but no allele exceeded 35% of the sequencing reads, then it was considered chimeric (Chi). **C)** A comparison of wild-type and edited DNA sequence of the homozygous 59 bp deletion in the male sterile line used in panel D. Highlighted in red is sgRNA1 used in pEM003 targeting exon 2. **D)** PCR amplicons spanning the target site in a wild-type plant or T_0_ lines homozygous or heterozygous for the 59 bp deletion were separated on a 3% agarose gel. The unlabeled lane has DNA size standards.

We generated two T-DNAs, each expressing two sgRNAs targeting either exon 2 or 3 and a *Cas9-P2A-TREX2* fusion under the *ZmUbi1* promoter (Chamness et al. 2023). TREX2, a 3’ exonuclease, increases both the size and frequency of insertion/deletion (indel) mutations created by Cas9, allowing easy detection of larger deletions by gel electrophoresis (Weiss et al. 2020; Čermák et al. 2017; Liu et al. 2024). Each construct also included the reporter genes *dsRED* (driven by the *S. viridis OLEOSIN4* promoter (*SvOle4*) and luciferase (driven by the *CmYLCV* promoter). The resulting constructs, pEM003 and pEM004, were transformed into *S. viridis* via Agrobacterium-mediated transformation (Nguyen et al. 2020).

Transformation yielded 43 plants – 23 with pEM003 and 20 with pEM004. PCR and sequencing revealed that 37 of the 43 T_0_ plants contained edits at either sgRNA1 or sgRNA2 (Figure 1B). Both constructs yielded T_0_ plants with homozygous inactivating mutations at individual sgRNA sites. One large in-frame deletion of 162 bp was observed for pEM004 but this T_0_ was not pursued further (Supplemental Figure 2). Notably, in pEM003 plant 21, a 59 bp deletion at sgRNA1 was recovered (Figure 1C). This deletion produced a clear mobility shift on a 3% agarose gel, enabling convenient genotyping (Figure 1D). Line 21 was selected for downstream phenotypic analyses. We noted that reporter gene expression varied among events and was not reliable for identifying transgenic individuals (Supplemental Figure 3).

### Identification and phenotypic characterization of male-sterile lines

To segregate away the T-DNA, line 21 and eight other T_0_ lines were backcrossed to wild-type ME034V. BC_1_F_1_ plants carried a variety of edits at the sgRNA1 and sgRNA2 sites (Supplemental Table 1). One BC_1_F_1_ individual derived from line 21, which lacked Cas9 as determined by PCR, was selfed to produce a BC_1_F_2_ population segregating for the 59 bp deletion.

Homozygous mutants exhibited a slightly shorter stature (Figure 2A) and smaller panicles (Figure 2B) relative to wild-type ME034V. Floral dissections revealed no differences in the number of anthers or stigmas, but at anthesis, line 21 anthers did not release pollen (Figure 2C-D). Potassium iodide (PI) staining confirmed the absence of pollen in line 21 anthers, validating the male-sterile phenotype (Figure 2E-H). No seeds were produced in the absence of outcrossing; however, seed set was restored when fertile pollen donors were used. All genotypes flowered approximately 18 days after planting under a 14-hour photoperiod in greenhouse conditions.

**Figure 2.**
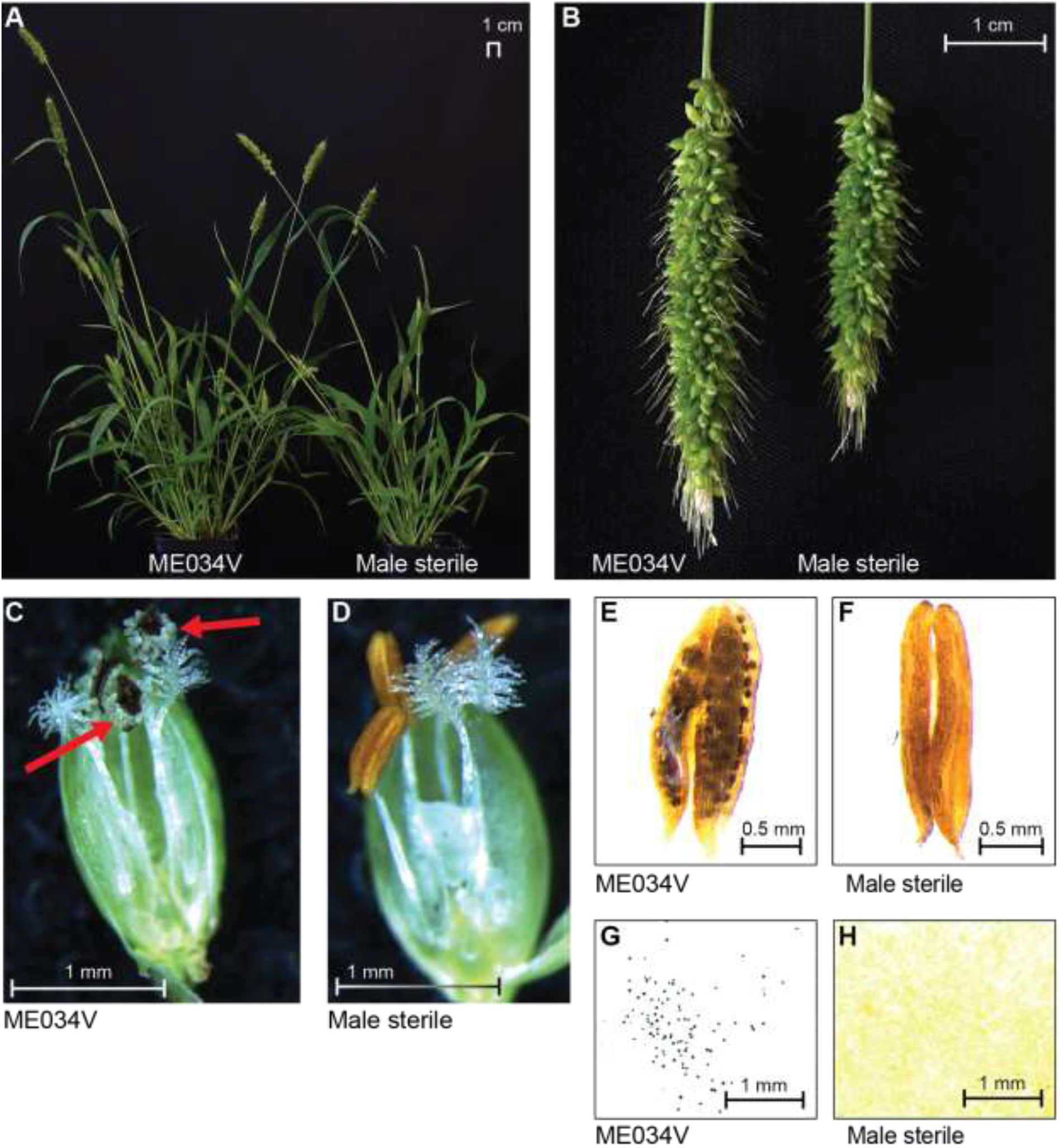
Phenotypic comparison of ME034V and male-sterile line 21. **A)** Visual comparison of ME034V and line 21 plants and **B)** Panicles. **C-D)** Flowers at anthesis. The red arrows indicate pollen. **E-F)** Anthers stained with potassium iodide. **G-H)** Pollen stained with potassium iodide.

### Development of a highly efficient crossing protocol

Using male-sterile line 21, we developed a simple and efficient crossing protocol (Supplemental Document 1). Seven to ten days after planting, progeny of heterozygous plants were genotyped to identify homozygous mutants for use as female parents. To prevent unintended pollination, panicles or entire plants were bagged at the time of panicle emergence. At anthesis, a panicle from a male parent of similar developmental stage was added to the bag, and the plants were placed in a crossing chamber with a pre-dawn cold treatment as described (Jiang et al. 2013). During the anthesis window, panicles were tapped every 15–20 minutes for two hours to facilitate pollination. This process was repeated over 2–3 consecutive days. Male donor panicles were removed 3–7 days after pollination, and the female panicles were re-bagged until seed harvest.

Crosses between male-sterile ME034V and A10.1, ME034V or AmilGFP donors yielded 49-145 F_1_ seeds per main panicle and 20 to 50 F_1_ per tiller panicle under manual pollination with tapping and cold treatment (Figure 3). Without tapping and cold treatments, only 20-32 seeds were produced per main panicle and 5-20 seeds per tiller panicle. Using two male donor panicles per female increased seed set to 34-62 seeds per main panicle.

**Figure 3.**
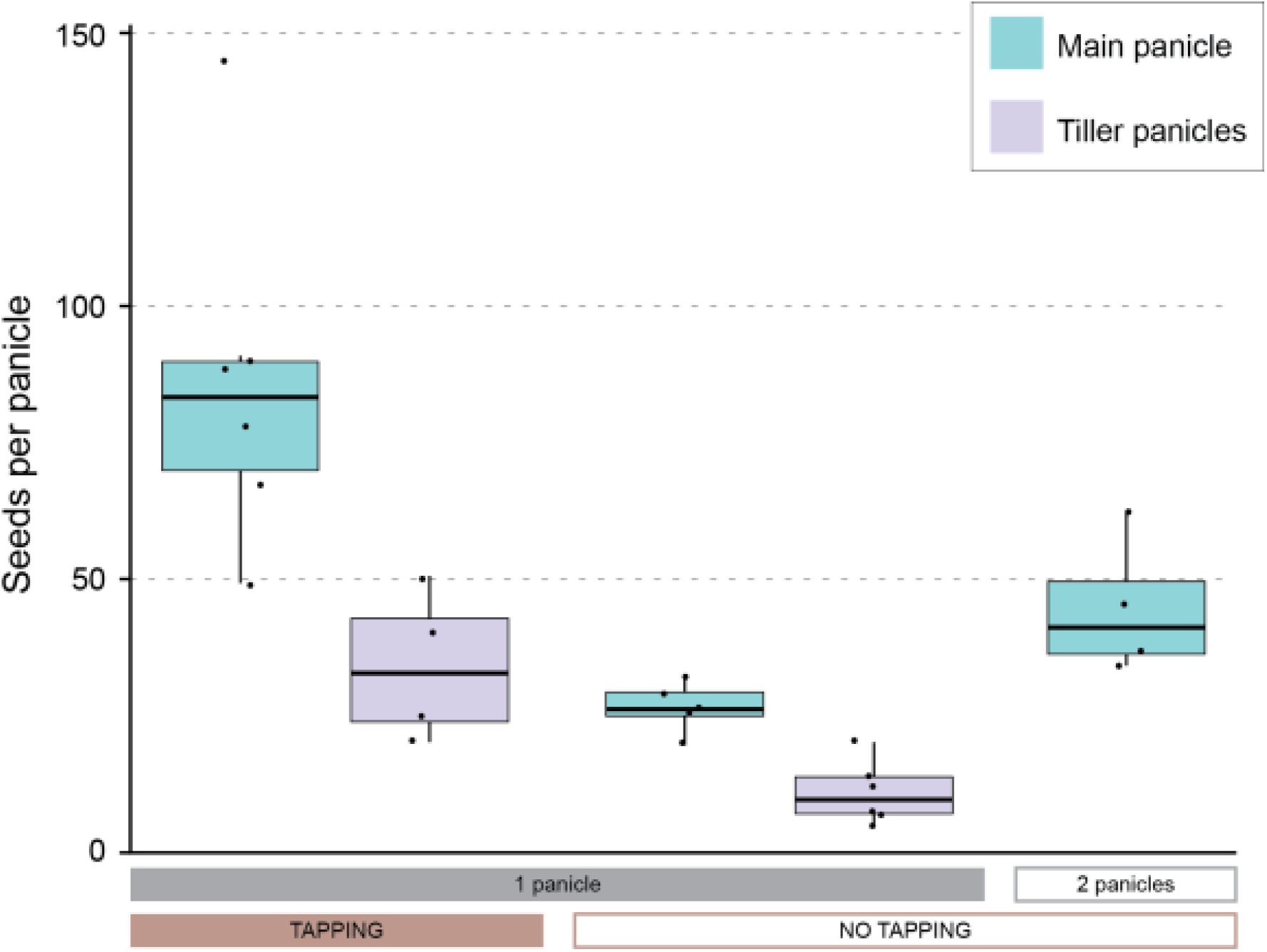
F_1_ seed production using different crossing strategies with the male-sterile line. A box plot illustrates F_1_ seed produced using various crossing strategies. Seed production in both the main panicles (teal) and tiller panicles (purple) of line 21 was compared. Crosses were conducted either through manual pollination (tapping) or in a greenhouse without manual pollination (no tapping). One group used two male panicles per cross, while all others used a single male panicle. The box plot represents the median, interquartile range, and variability among individual plants.

### Purifying genetic backgrounds of male sterile lines

To eliminate background mutations introduced during transformation, we began backcrossing pEM003_21_BC_1_F_2__31 to wild-type ME034V. Our goal is to develop a BC_8_F_2_ population with >99% ME034V background. BC_4_F_1_ seeds have been harvested, and backcrossing is ongoing (Supplemental Figure 4).

Because NMU-mutagenized populations are in the A10.1 background (Huang et al. 2017), we also developed a Cas9-free male-sterile line in A10.1 using the pEM677 construct. This construct was similar to pEM003, except that it contained *ZmUbi1* driving *AmilGFP* in place of the *dsRed* and luciferase reporters. Line pEM677_1.18.24 was identified as Cas9-free and homozygous for 61 bp deletions at both sgRNA1 and sgRNA2 sites, distinguishable by PCR on a 3% agarose gel (Supplemental Figure 5). Backcrossing to wild-type A10.1 is in progress and we are currently growing BC_1_F_1_ plants.

### Estimating outcrossing frequencies in greenhouse conditions

To assess pollen flow, seeds harvested from heterozygous plants were planted in a tray and genotyped at 10d. Panicles of homozygous male-sterile plants were bagged either before or after anthesis (Supplemental Table 2). No seeds developed when bagging occurred before anthesis, but 2-30 seeds were recovered when bagging occurred after anthesis initiation, indicating the potential for unintended outcrossing with their heterozygous or wild-type siblings.

To quantify outcrossing, wild-type ME034V and AmilGFP plants were grown in adjacent trays spaced 15–20 cm apart (Figure 4A). ME034V panicles were bagged after flowering, and seeds were harvested from main panicles at five weeks and from tiller panicles at seven weeks after planting. After seed coat removal, seeds were screened for GFP fluorescence (Figure 4B), and the percentage of GFP-positive seeds was calculated per plant.

**Figure 4.**
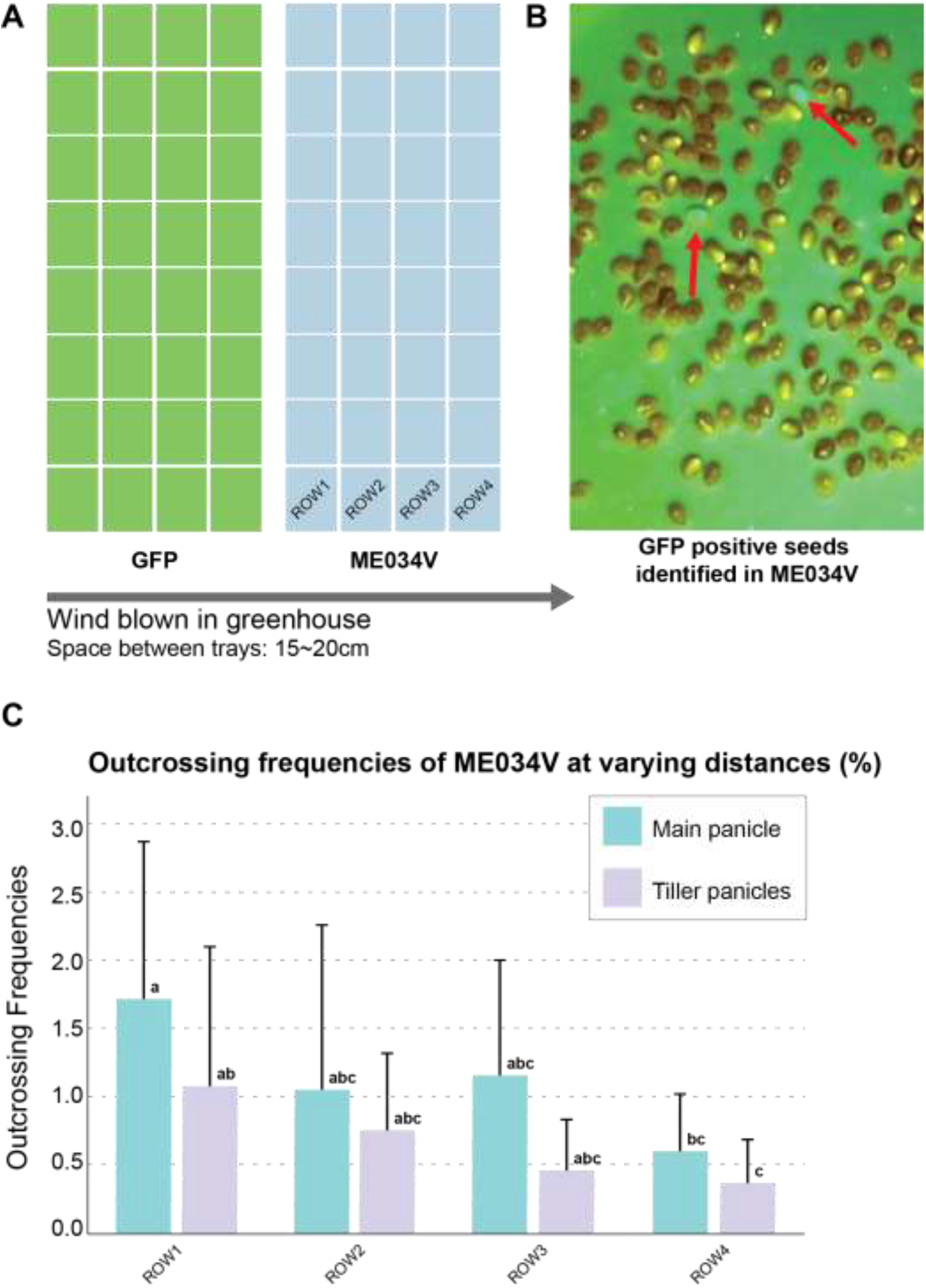
Outcrossing frequencies of wild-type ME034V under greenhouse conditions. **A)** Experimental design of outcrossing frequency tests between trays in the greenhouse. **B)** Red arrows indicate GFP-positive seeds identified from a pool of seeds harvested from the panicles of ME034V. **C)** Outcrossing frequencies of main and tiller panicles of ME034V in the greenhouse. The figure shows significant variation both between and within rows.

Outcrossing frequencies were compared across four rows of ME034V plants at increasing distances from the *AmilGFP* donor tray. Frequencies ranged from 0.6% to 3.5% in main panicles and 0.2% to 3.1% in tiller panicles (Figure 4C). Row 1, closest to the donor tray, exhibited the highest average outcrossing frequencies (1.7% ± 1.1 in main panicles and 1.1% ± 1.0 in tiller panicles), while Row 4 had the lowest (0.6% ± 0.4 in main panicles, 0.4% ± 0.3 in tiller panicles). When comparing outcrossing frequencies between main and tiller panicles within the same row, a significant difference (p < 0.05) was observed only in Row 4, with no significant differences in the other rows.

## Discussion

We successfully generated male-sterile lines in the commonly used *Setaria viridis* accessions ME034V and A10.1 by targeting the *NO POLLEN 1* (*NP1*) ortholog using CRISPR/Cas9. The resulting alleles are small deletions readily distinguishable by PCR, enabling rapid identification of homozygous mutants for crossing and heterozygotes for maintaining the line. These resources address a key limitation in *S. viridis* genetics: the labor-intensive and technically challenging process of manual crossing.

Using these male-sterile lines, we developed a simple, scalable, and highly efficient crossing protocol that eliminates the need for emasculation. Traditional methods relied on heat treatments, trimming, and precise manual pollination, all of which required specialized training and yielded inconsistent results (Jiang et al. 2013). Our revised approach—based on co-bagging male and female panicles—reduces labor dramatically: a single researcher can process over 100 panicles per day, compared to only ∼15 using the previous method. Critically, the protocol requires minimal equipment and is adaptable to both greenhouse and field conditions, including environments where cold-treatment chambers are unavailable.

Beyond efficiency, the new protocol improves reliability and reproducibility. F_1_ seed yield per panicle has increased significantly (20–32 seeds without tapping panicles together; up to 145 seeds with tapping panicles together and cold treatment), and hybrid purity is guaranteed by the absence of functional pollen in the female parent. This contrasts sharply with previous protocols, where high rates of self-pollination led to contamination and reduced confidence in genetic experiments.

To ensure genetic integrity, we have initiated backcrossing of both ME034V and A10.1 male-sterile lines to their respective wild-type backgrounds. Our goal is to remove any background mutations introduced during transformation or tissue culture by completing eight rounds of backcrossing. However, seed is available for community use prior to the completion of this process.

The ME034V male-sterile line also enabled us to estimate the pollen flow under greenhouse conditions, and led us to investigate the outcrossing rate in Setaria using wild-type and the AmilGFP line. Observed outcrossing frequencies (0.6% to 3.5%) are consistent with previous estimates for *Setaria* under natural conditions (Till-Bottraud et al. 1992; Li et al. 1945). Importantly, our results underscore the need to bag panicles immediately after emergence before anthesis to prevent unintended pollination. Given the close spacing and overlapping inflorescences in greenhouse settings, we suspect that most pollen transfer occurs via direct floral contact rather than airborne pollen flow.

In summary, the development of male-sterile lines and a robust crossing protocol represents a major advancement for *S. viridis* as a model system. These tools will facilitate efficient generation of F_1_ hybrids, accelerate development of recombinant inbred lines (RILs) and multi-parent advanced generation intercross (MAGIC) populations, and streamline approaches such as bulked segregant analysis. More broadly, the availability of male-sterile lines enhances the scalability of forward genetics and functional genomics studies in *S. viridis*, supporting its continued role as a premier model for C_4_ plant biology.

## Materials and Methods

### Plant materials, growth conditions and *S. viridis* transformation

*S. viridis* accession ME034V served as the male parent for backcrossing. Accession A10.1 was used for assessing crossing efficiency. T_3_ seeds from a *S. viridis* AmilGFP line #1-9-2 were used to quantify outcrossing frequencies. Plants were cultivated either in a greenhouse or a high-light growth chamber. Greenhouse conditions were 31°C/22°C (day/night) with a 14 h light/10 h dark photoperiod and 40% relative humidity. Growth chamber conditions were 31°C/22°C (day/night) under a 16 h light/8 h dark photoperiod with CO_2_ concentration maintained at approximately 19,000 ppm, 50% relative humidity, and light intensity of 400 µmol m^−2^ s^−1^. Controlled crosses conducted in the chamber were preceded by a pre-dawn cold treatment at 15°C from 07:00 to 07:30 AM.

Plasmid constructs pEM003, pEM004 and pEM677 were introduced into *Agrobacterium tumefaciens* strain AGL1 and transferred into calli of *S. viridis* ME034V or A10.1 via Agrobacterium-mediated transformation at the Donald Danforth Plant Science Center Tissue Culture Facility. T_0_ plants were transplanted into soil and grown either in the greenhouse for seed harvest or in the growth chamber for use in crossing experiments.

### Backcrossing and phenotypic characterization of the male-sterile ME034V line

To eliminate background mutations introduced by CRISPR/Cas9 or tissue culture, male-sterile T_0_ plants were backcrossed to ME034V using the anther-to-stigma method under a dissecting microscope (Jiang et al. 2013). BC_1_F_1_ seeds were harvested two weeks after pollination and dried at 30°C. Dormancy was broken by soaking seeds in 5% liquid smoke for 20–24 hours at room temperature while enclosed in mesh biopsy bags (AutoSette Series, 30 × 50 mm) (Sebastian et al. 2014). Bags were briefly dried in a fume hood for 30-60 min and then placed in moist, long-fiber sphagnum moss within sealed plastic bags and stratified at 4°C for two weeks. Stratified seeds were either sown directly or dried at 30°C for 1–2 days prior to sowing or storage. BC_1_F_1_ seeds were grown and self-pollinated to produce BC_1_F_2_ seeds. These plants were genotyped and homozygous male-sterile individuals were selected for additional backcrossing to ME034V using the crossing protocol developed in this study. This backcrossing process is being repeated to obtain BC_8_F_2_ seeds, which are expected to carry >99% of the ME034V background. Seeds of the male-sterile lines are available to the Setaria research community upon request.

Whole plants and panicles were photographed using a Nikon D5100 digital camera. Flowers were dissected and imaged using a Leica M125C stereomicroscope. Anthers and pollen grains were stained with iodine–potassium iodide (I_2_-KI) solution consisting of 0.2% iodine and 2% potassium iodide and observed under the stereomicroscope.

### DNA extraction and genotyping

Genomic DNA was extracted from plant tissues using a modified CTAB method (Fulton et al. 1995) or the Phire Plant Direct PCR Master Mix (Thermo Scientific, Cat. No. F160L), according to the manufacturer’s instructions. To detect the presence of the Hygromycin phosphotransferase (*hpt)* region in transgene, a duplex PCR was performed using primer pairs hpt_F/R and *Sv*_F/R (Supplemental Table 3), amplifying 335 bp and 540 bp fragments from the *hpt* and *Setaria romosa 1* (*Sv*.) genes, respectively, with the latter serving as an internal control. The 59 bp deletion in *SvNP1* was detected using primers MS_F and MS_R (Supplemental Table 3). For CTAB-extracted DNA, PCR reactions were carried out using GoTaq® Master Mix (Promega, Cat. #M7123) in 25 µL volumes containing 12.5 µL of 2× mix, 1 µL of each primer, 2 µL of DNA (10–50 ng/µL), and water. Thermal cycling conditions for *hpt* and *Sv*. gene detection were: 95°C for 3 minutes; 28 cycles of 95°C for 15 seconds, 60°C for 15 seconds, and 72°C for 20 seconds; final extension at 72°C for 10 minutes. For the 59 bp deletion, PCR cycling included 40 cycles with an annealing temperature of 66.4°C and extension at 72°C for 30 seconds.

For reactions using Phire Plant Direct Master Mix, the PCR mixture contained 12.5 µL of 2× Phire Direct Buffer, 1 µL of each primer, 2 µL of DNA extract, and water to 25 µL. Thermal cycling for hpt and Sv. gene detection included an initial denaturation at 98°C for 5 minutes, followed by 28 cycles at 98°C for 15 seconds, 66.8°C for 15 seconds, and 72°C for 20 seconds, with a final extension at 72°C for 10 minutes. For the 59 bp deletion, cycling consisted of 40 cycles of 98°C for 15 seconds, with annealing and extension combined at 72°C for 45 seconds.

PCR products were separated on 3% agarose gels run at 100 V for 45 minutes and visualized using either a 100 bp or low molecular weight ladder (New England Biolabs). For sequencing, PCR products were gel-purified using the DNA Clean & Concentrator-5 kit (Zymo Research) and submitted to Genewiz for Sanger sequencing. Sequence data were analyzed using Benchling and Synthego ICE software.

### Measuring outcrossing frequencies

Wild-type ME034V and AmilGFP plants were grown in adjacent 4×9 cell trays with 20 cm spacing. ME034V main and tiller panicles were bagged at four and five weeks after planting, respectively. Seeds were harvested two weeks after bagging, and seed coats were removed using a custom-built *Setaria* dehuller (Van Eck & Swartwood 2015). Dehulled seeds were screened for GFP fluorescence using the NIGHTSEA Xite™ Fluorescence Flashlight System (Xite-CY) and barrier filter glasses (FG-CY).

### T-DNA design and cloning

T-DNA constructs were assembled using Golden Gate cloning (Čermák et al. 2017). sgRNAs targeting *Sevir*.*9G353400* were designed using CRISPOR (https://crispor.org) and selected based on low off-target scores and predicted overlap with conserved protein domains identified through Phytozome (https://phytozome-next.jgi.doe.gov/blast-search). sgRNA fragments were amplified using Q5 polymerase (New England Biolabs), gel-purified, and cloned into pMOD_B2520 using BsaI, generating intermediate constructs pEM001 and pEM002. These were combined with additional modules to form the final T-DNA constructs pEM003 and pEM004. These included pMOD_A1910 containing ZmUbi::TREX2-P2A-Cas9, pMOD_C’_SvOle4::dsRed, and pMOD_D_CmYLCV::FLUC, with final assembly performed using AarI digestion (Chamness et al. 2023). Primers and plasmids used in this study are listed in Supplemental Table 3 and Supplemental Table 4, respectively.

### Genotyping and analysis of heritable mutagenesis

Genomic DNA from T_0_ plants was used to amplify target regions using primers located at least 160 bp outside the sgRNA sites (EM407/408 or EM445/446, Supplemental Table 3). PCR products were sequenced (Eurofins) and analyzed using Synthego ICE (Conant et al. 2022). Mutation outcomes were categorized as homozygous if a single sequence accounted for >85% of the reads, heterozygous if one or two alleles each comprised 35–65%, or chimeric if multiple alleles each comprised >10%. Once line pEM003-21 was identified as a homozygous 59 bp deletion mutant, new primers (MS_F/MS_R, Supplemental Table 3) were designed to produce PCR products of 255 bp in edited lines and 314 bp in wild-type, with both bands present in heterozygotes. PCR was conducted using an annealing temperature of 66.4°C for 15 seconds and an extension of 30 seconds. Products were separated on 3% agarose gels run at 100 V for 45 minutes.

### Generation of an *AmilGFP*-expressing *Setaria* line

A *Zea mays* codon-optimized *Acropora millepora* green fluorescent protein (*ZmCoAmilGFP*) sequence was synthesized and cloned into a Golden Gate Level 0 (L0) module (Liljeruhm et al. 2018), (Weber et al. 2011). For constitutive expression, the *ZmUbi1* promoter was used to drive *AmilGFP* expression. The *pZmUbi1::AmilGFP* L1 transcriptional unit was assembled with a hygromycin expression L1 cassette to generate a L2 binary vector (Supplemental Figure 6). This binary vector was transformed into *S. viridis* as described above. Transgenic lines were screened for *GFP* fluorescence using a SpectraMax® M3 plate reader (Molecular Devices, San Jose, USA) set to 485 nm excitation and 515 nm emission wavelength. *GFP* fluorescence co-segregated with the presence of *hpt* and was stably inherited across multiple generations. Line #1-9-2 (T_3_), which displayed strong *GFP* fluorescence, was used as a pollen donor in outcrossing experiments.

## Acknowledgements

We thank Tuany Volz for assistance with the outcrossing frequency experiments, Collin Luebbert for help with statistical analysis and box plot generation, and Kit Leffler for support with formatting, figures, and tables.

## Author Contributions

HJ, EM, DV and IB conceived and designed the research, HJ, EM, BM and DL, conducted the experiments, H.J., HJ, EM, BM, DV and IB wrote and edited the manuscript, DN reviewed and edited the manuscript.

