## Supplemental Figure 1 for "Development of male-sterile lines of *Setaria viridis* to accelerate C_4_ model plant genetics"

**Supplemental Figures**

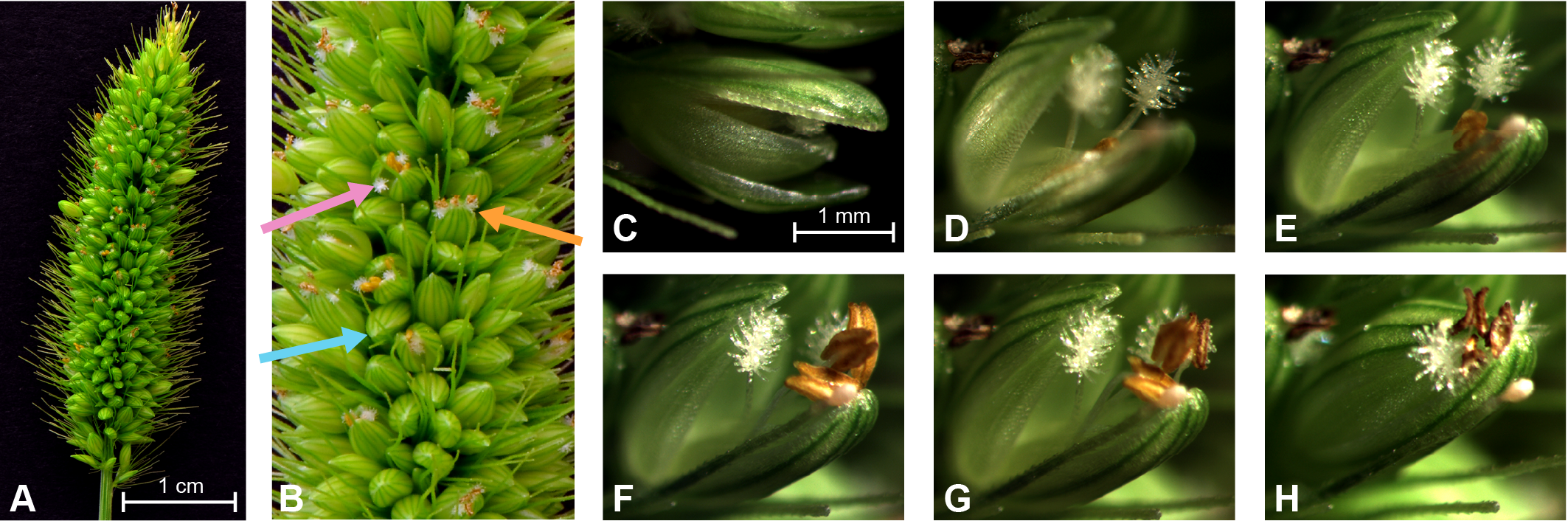

**Supplemental Figure 1.** Flower progression in *S. viridis* A10.1 during anthesis (~40-min. total duration) **A)** *Setaria* panicle at anthesis. **B)** Enlarged view of the middle section of the panicle. Arrows indicate flowers at different stages: before opening (blue, bottom arrow on the left), opening (orange, arrow on the right), and blossomed (pink, top arrow on the left). **C–H)** Sequential progression of a flower from opening to closing over the course of ~40 minutes.
