## Supplemental Figure 2 for "Development of male-sterile lines of *Setaria viridis* to accelerate C_4_ model plant genetics"

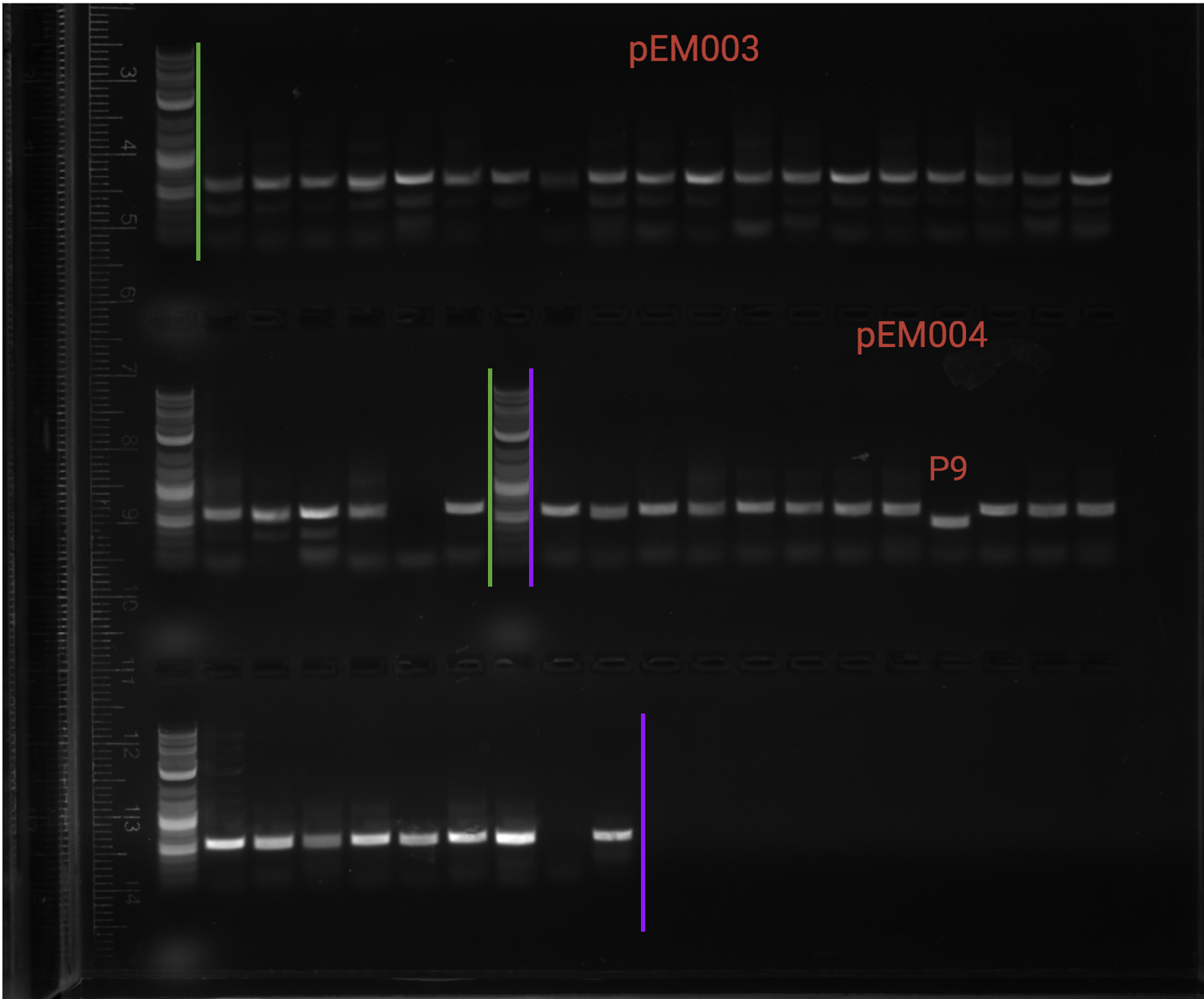

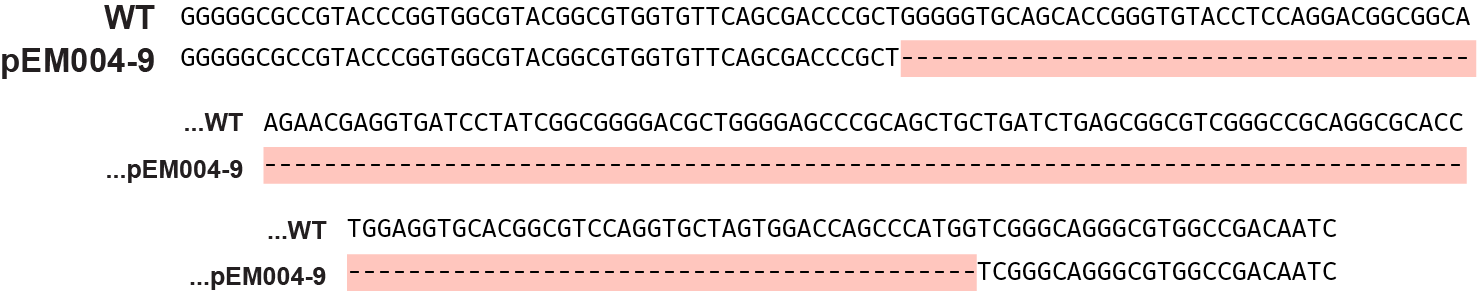

**Supplemental Figure 2.** PCR products used for genotyping the T_0_ generation. The vertical green lines delimit progeny derived from pEM003; the vertical purple lines delimit progeny derived from pEM004. Plant 9 of pEM004 had a large homozygous deletion of 162 bp between both sgRNAs. The DNA sequence of this deletion is shown below the gel.
