## Supplemental Figure 3 for "Development of male-sterile lines of *Setaria viridis* to accelerate C_4_ model plant genetics"

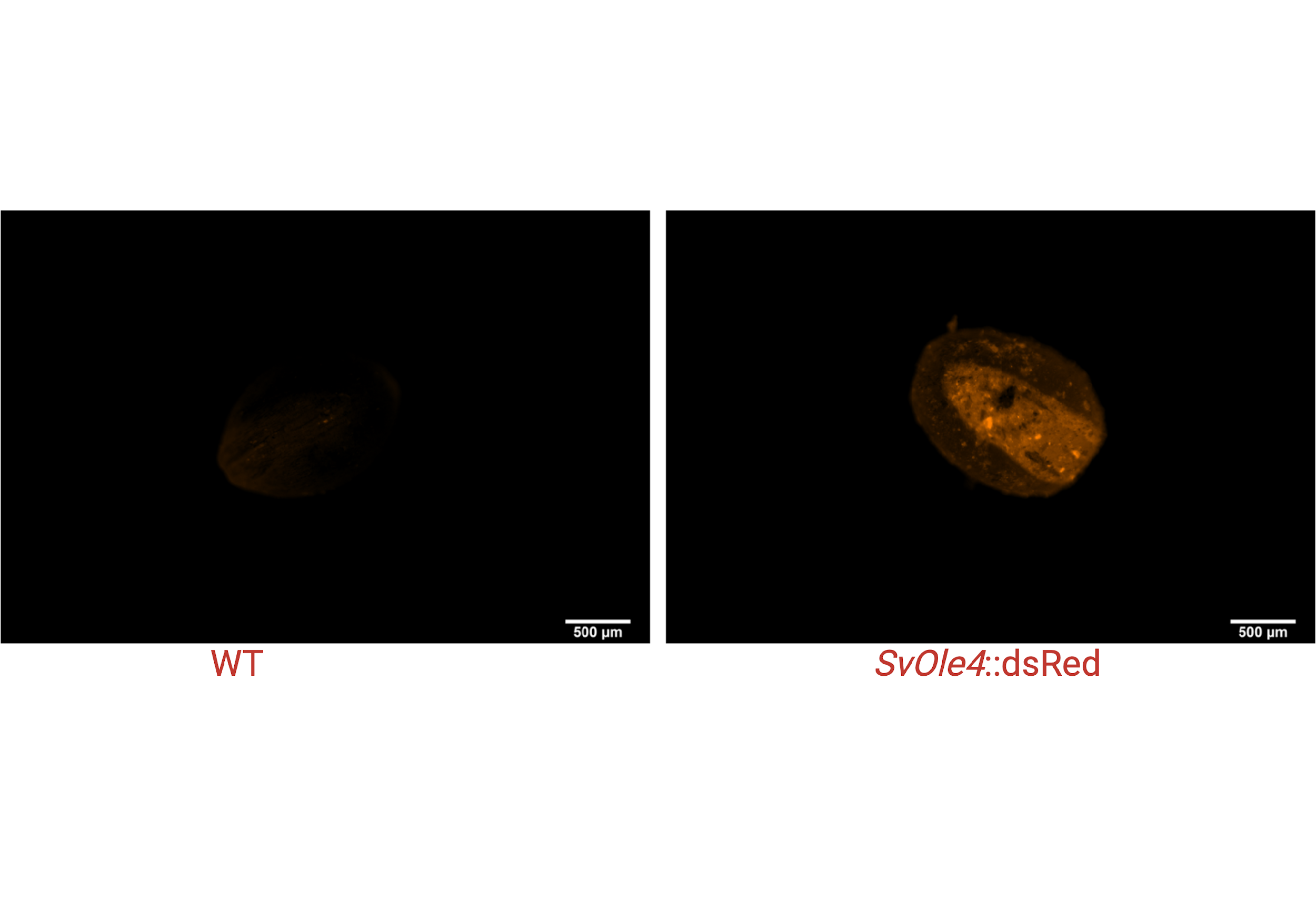

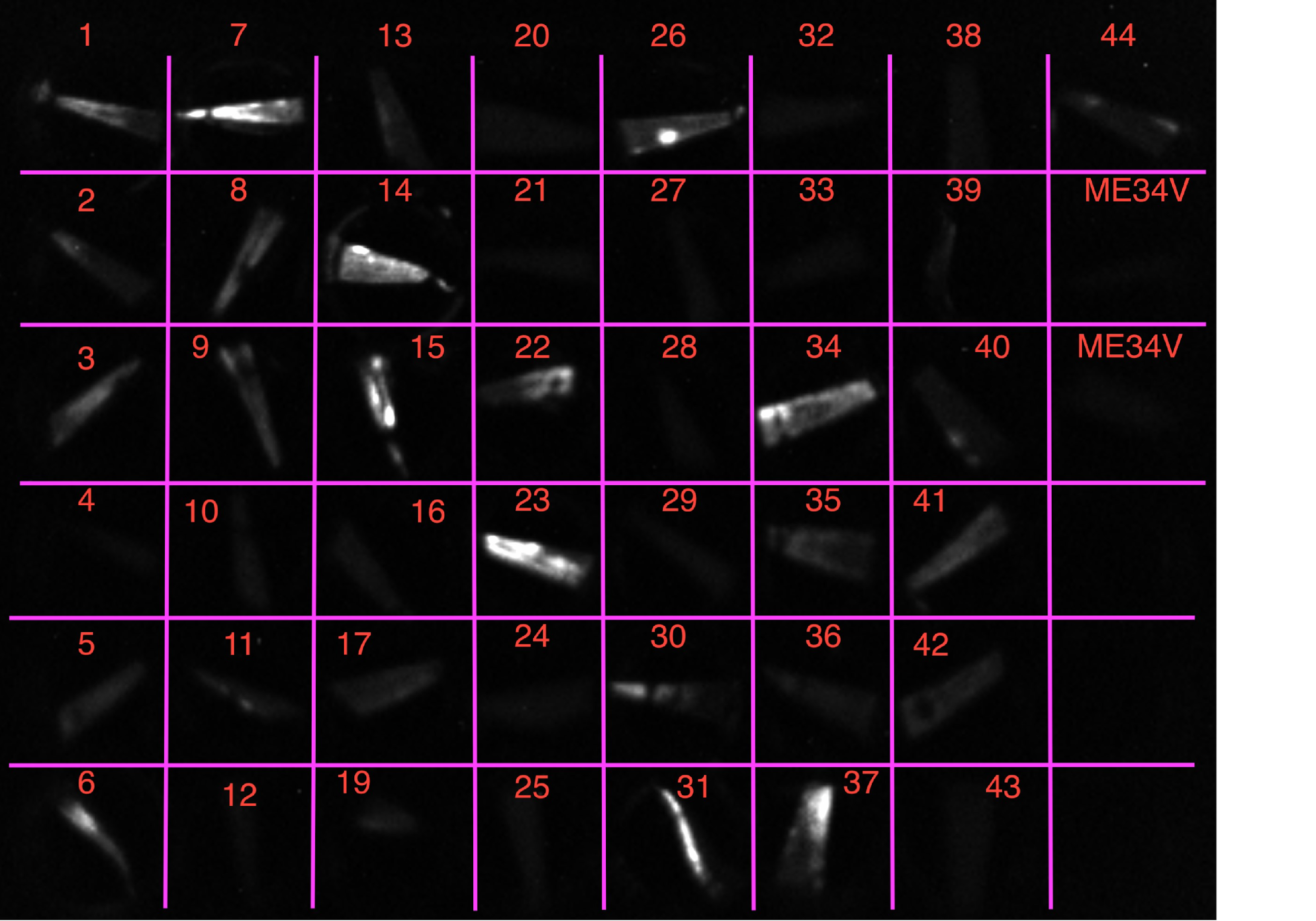

**Supplemental Figure 3.** The upper right panel is a representative photo of a T_0_ *S. viridis* line expressing dsRed in the seed using the *SvOle4* promoter. A wild-type seed under the same illumination is shown in the upper left panel. The lower panel shows a luciferase assay on excised leaves from all T_0_ plants.
