## Supplemental Figure 4 for "Development of male-sterile lines of *Setaria viridis* to accelerate C_4_ model plant genetics"

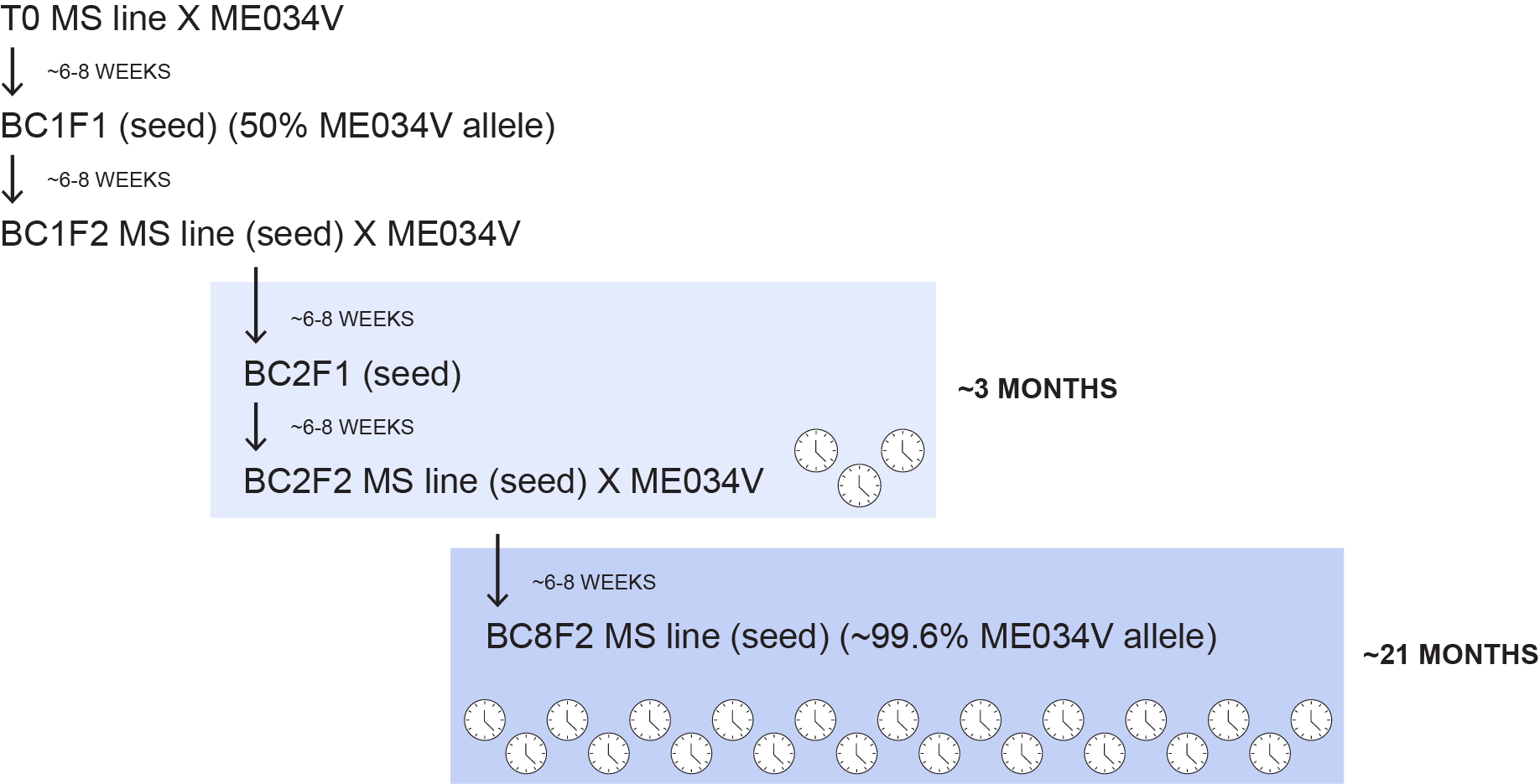

**Supplemental Figure 4.** Schematic of the backcrossing process with ME034V and the required timeframe.
