## Supplemental Figure 5 for "Development of male-sterile lines of *Setaria viridis* to accelerate C_4_ model plant genetics"

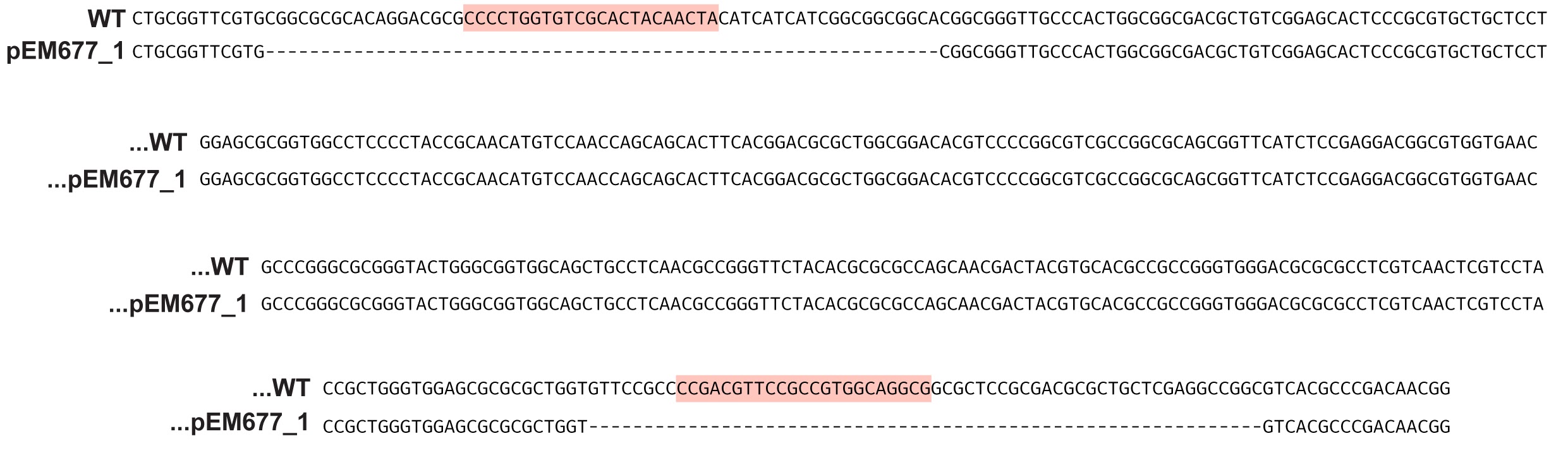

**Supplemental Figure 5.** A DNA sequence alignment of the homozygous 61 bp deletions observed at both sgRNA1 and sgRNA2 targeting exon 2 in the male sterile line in the A10.1 background.
