## Supplemental Figure 6 for "Development of male-sterile lines of *Setaria viridis* to accelerate C_4_ model plant genetics"

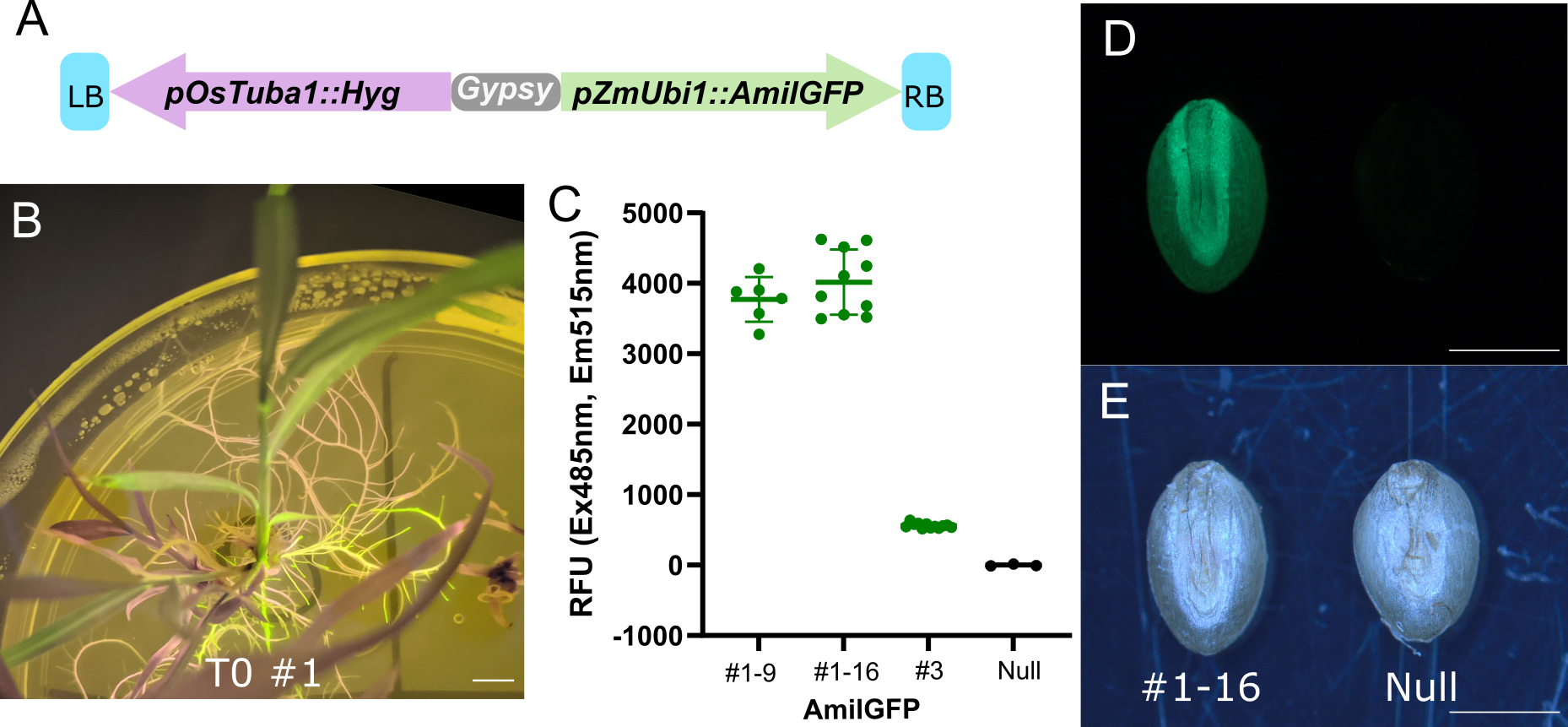

**Supplemental Figure 6. AmilGFP overexpression and characterization*.* A)** A schematic for the transformation construct used to express AmilGFP (pZmUbi1::AmilGFP). **B)** GFP fluorescence of a primary *pZmUbi1::AmilGFP* transformant. GFP fluorescence (green) and chlorophyll autofluorescence (pink) are observed with imaging through an amber plate on top of a blue-light gel transilluminator. Scale bar = 5mm. **C)** GFP variation from individual AmilGFP transgenic events. Fluorescence was scored from leaf punches from individual T_2_ lines on a plate reader with the indicated excitation and emission spectra, #1-9 and #1-16 are sibling T_2_ lines carrying multi-copy T-DNA insertions, #3 is a single insertion, and null is a transgene-free segregant sibling from #3. **D-E)** GFP image of the embryo side of the AmilGFP seed (left) and null (right). GFP image (D) and bright-field image (E) of the embryo side of *S. viridis* seeds. Scale bar = 1 mm.
