## Supplemental Table 1 for "Development of male-sterile lines of *Setaria viridis* to accelerate C_4_ model plant genetics"

**Supplemental Tables**

| **T_0_ family** | **BC_1_F_1_ Plant ID** | **Hpt** | **gRNA1** | **gRNA2** |
| --- | --- | --- | --- | --- |
| pEM003_4 | F1_2 | + | 2 bp | 1 bp del |
|  | F1_3 | - | Edits inconclusive; potential large deletion | 3 bp del |
| pEM003_6 | F1_4 | + | 7 bp | 1 bp del |
| pEM003_8 | F1_6 | + | 1 bp insert | 19 bp del |
| pEM003_16 | F1_17 | + | 5 bp | 1 bp del |
| pEM003_21 | F1_29 | + | Edits inconclusive; potential large deletion | 2 bp del |
|  | F1_30 | - | Edits inconclusive; potential large deletion | 2 bp del |
|  | F1_31 | - | Edits inconclusive; potential large deletion | 2 bp del |
|  | F1_32 | + | Edits inconclusive; potential large deletion | 2 bp del |
| pEM003_18 | F1_33 | - | 13 bp | WT |
| pEM003_19 | F1_34 | + | 2 bp | 14 bp del |
| pEM004_3 | F1_2 | + | 5 bp | 4 bp del |
|  | F1_3 | + | 5 bp | 4 bp del |
|  | F1_4 | - | 3 bp | 28 bp del |
| pEM004_17 | F1_10 | + | 14 bp | 11 bp del |
|  | F1_11 | + | 4 bp | 6 bp del |
|  | F1_12 | + | 1 bp | 6 bp del |

**Supplemental Table 1:** Edits identified in BC_1_F_1_ plants derived from the backcross of male sterile T_0_ plants with the ME034V parent.
