## Supplemental Table 2 for "Development of male-sterile lines of *Setaria viridis* to accelerate C_4_ model plant genetics"

| **Plant ID** | **Time to Bag Panicles** | **# of Panicles Bagged/plant** | **# of Panicles with Seed** | **Seed Number** |
| --- | --- | --- | --- | --- |
| ME034V | Before anthesis | 1 | 1 | 300-500 |
| ME034V | After anthesis | 1 | 1 | 300-500 |
| 3.4 | Before anthesis | 1 | 0 | 0 |
| 3.17 | Before anthesis | 1 | 0 | 0 |
| 3.23 | Before anthesis | 3 | 0 | 0 |
| 31.1 | Before anthesis | 3 | 0 | 0 |
| 31.2 | Before anthesis | 3 | 0 | 0 |
| 31.3 | Before anthesis | 3 | 0 | 0 |
| 31.5 | Before anthesis | 3 | 0 | 0 |
| 31.13 | Before anthesis | 1 | 0 | 0 |
| 3.3 | ~1d after anthesis initiation | 2 | 1 | 3 |
| 3.8 | ~1d after anthesis initiation | 1 | 1 | 2 |
| 31.13 | ~1d after anthesis initiation | 1 | 1 | 10 |
| 31.1 | ~10 d after anthesis initiation | 1 | 1 | ~30 |
| 31.2 | ~10 d after anthesis initiation | 1 | 1 | ~20 |

**Supplemental Table 2**: Comparison of panicle bagging times and seed set
