## Supplemental Table 3 for "Development of male-sterile lines of *Setaria viridis* to accelerate C_4_ model plant genetics"

| **Oligo Name** | **DNA sequence (5’ to 3’)** | **Purpose** |
| --- | --- | --- |
| EM001 | CACGTCTCGTTGTGTAGTTGTAGTGCGACACCAGGTTTTAGAGCTAGAAATAGCAAG | Used to clone sgRNA1 into pMOD_B2520 for pEM003 |
| EM002 | GTCGTCTCCGGAGAAAAAAAGCACCGACTCGGTGCCACT |  |
| EM003 | CACGTCTCGCTCCGGGATCATGAACCAACGGCCT | Used to clone sgRNA2 into pMOD_B2520 for pEM003 |
| EM004 | GTCGTCTCCAAACACGTTCCGCCGTGGCAGGCGCACAAGCGACAGCGCGCGGGTTTA |  |
| EM005 | CACGTCTCGTTGTGACCCGGTGCTGCACCCCCAGGTTTTAGAGCTAGAAATAGCAAG | Used to clone sgRNA1 into pMOD_B2520 for pEM004 |
| EM002 | GTCGTCTCCGGAGAAAAAAAGCACCGACTCGGTGCCACT |  |
| EM003 | CACGTCTCGCTCCGGGATCATGAACCAACGGCCT | Used to clone sgRNA2 into pMOD_B2520 for pEM004 |
| EM006 | GTCGTCTCCAAACACCATGGGCTGGTCCACTAGCACAAGCGACAGCGCGCGGGTTTA |  |
| MS_F | GCCTCGTCGCACTCTCAGAGGA | Used for quick screening of the 59 bp deletion on 3% agarose gels |
| MS_R | GCGTCCGTGAAGTGCTGCTGGTT |  |
| EM407 | GCTCCTTGTGCCTCGTCGCA | Used for genotyping of edits in exon 3 of Sevir.9G353400 |
| EM408 | GCAGCACGCGGGAGTGCTCC |  |
| EM445 | CCGACTTCCTCCGCCACGC | Used for genotyping of edits in exon 4 of Sevir.9G353400 |
| EM446 | AGCTGCCCGGTCTGCAACTA |  |
| *hpt_F* | AGGCTCTCGATGAGCTGATGCTTT | Used for genotyping hygromycin resistance gene |
| *hpt_R* | AGCTGCATCATCGAAATTGCCGTC |  |
| *Sv_F* | CAGCAAGCCGCCTATATGGAG | Use for genotyping Setaria Romosa 1 gene |
| *Sv_R* | TCGTCTCAGGAGTGGCCAAGT |  |

**Supplemental Table 3.** Oligos used in this study.
