## Supplemental Table 4 for "Development of male-sterile lines of *Setaria viridis* to accelerate C_4_ model plant genetics"

| **Plasmid** | **Description** | **Level** | **Sequence Link** |
| --- | --- | --- | --- |
| pMOD_A1910 | Level 1 A position module containing ZmUbi1::TREX2-P2A-Cas9::AtHSP | 1 | [Sequence Link 1](https://benchling.com/s/seq-LPZD6JplQ1KOEAkA6VPg?m=slm-K4tBVB3IxD0W3FYfkCyo) |
| pMOD_B2520 | Level 1 B position destination module containing OsU6::ccdB::sgRNA scaffold | 1 | [Sequence Link 2](https://benchling.com/s/seq-FsM1mdWphXNIi0Xhq4qB?m=slm-fsBISBKYKU2ZYqWnLIHI) |
| pMOD_C'-pOle-dsRED-rbcS | Level 1 C' position module containing SvOle4::dsRed::Pea-rbcS | 1 | [Sequence Link 3](https://benchling.com/s/seq-nxBIbzVmL2v67sF7xEAe?m=slm-y3nZU4F7Mlh8JQWpNiFP) |
| pMOD_D-CmYLCV-Fluc-35S | Level 1 D position module containing CmYLCV::Fluc::35S | 1 | [Sequence Link 4](https://benchling.com/s/seq-9kkMdmsF1FVLAc3xF1qs?m=slm-PNQVXb12uGToeI2KLmeu) |
| pEM001 | Level 1 B position containing 2 sgRNAs targeting exon 3 of Sevir.9G353400 | 1 | [Sequence Link 5](https://benchling.com/s/seq-fsTU3TSv7WgcPPXVl7bC?m=slm-4PxkSfUcKQqYHP7oc00S) |
| pEM002 | Level 1 B position containing 2 sgRNAs targeting exon 4 of Sevir.9G353400 | 1 | [Sequence Link 6](https://benchling.com/s/seq-gIohlXtKo39zDS9yUfqj?m=slm-MdYMM1swnCXpOelESOTA) |
| pEM003 | T-DNA used for mutagenesis and targeting exon 3 of Sevir.9G353400 in ME034V | 2 | [Sequence Link 7](https://benchling.com/s/seq-wIZxbrpZ09msdAkjI9H2?m=slm-0K7ft3KqWL7gN7cUSzwe) |
| pEM004 | T-DNA used for mutagenesis and targeting exon 4 of Sevir.9G353400 in ME034V | 2 | [Sequence Link 8](https://benchling.com/s/seq-yhcSvUUdIIuZb6iampoV?m=slm-kXneyfdSmAsNgw0PZyNB) |
| pEM658 | Level 1 D position containing *ZmUbi1::AmilGFP* | 1 | [Sequence Link 9](https://benchling.com/s/seq-rw9gEf0rCSK09wkqPIAb?m=slm-aJQ6j9cBN5gjBmdb0npL) |
| pEM677 | T-DNA used for mutagenesis and targeting exon 3 of Sevir.9G353400 in A10 | 2 | [Sequence Link 10](https://benchling.com/s/seq-iYbrXg8CSDRV8PRoOvFe?m=slm-eapBSdVfJoFsvFDv8FAu) |
| pTRANS_250d | Level 2 VSI destination backbone containing Hygromycin resistance | 2 | [Sequence Link 11](https://benchling.com/s/seq-OU4CvdwcG9ZPvPjc7QUY?m=slm-mzaIoCsvzyFD5ZP5fGQN) |

**Supplemental Table 4.** Plasmids used in this study.
